# B-1 cell-mediated modulation of m1-macrophage profile can ameliorate microbicidal functions and disrupt the evasion mechanisms of *Encephalitozoon cuniculi*

**DOI:** 10.1101/567404

**Authors:** Adriano Pereira, Anuska Marcelino Alvares-Saraiva, Fabiana Toshie de Camargo Konno, Diva Denelle Spadacci-Morena, Elizabeth Cristina Perez, Mario Mariano, Maria Anete Lallo

## Abstract

Here, we have investigated the possible effect of B-1 cells on the activity of peritoneal macrophages in *E. cuniculi* infection. In the presence of B-1 cells, peritoneal macrophages had an M1 profile with showed increased phagocytic capacity and index, associated with the intense microbicidal activity, increased proinflammatory cytokines production and a higher percentage of apoptotic death. The absence of B-1 cells was associated with a predominance of the M2 macrophages, indicating reduced phagocytic capacity and index, microbicidal activity, proinflammatory cytokine production, and apoptotic death, but equal death rate. In addition, in the M2 macrophages, spores of phagocytic *E. cuniculi* with polar tubular extrusion were observed, which is an important mechanism of evasion of the immune response. The results showed the importance of B-1 cells in the modulation of macrophage function against *E. cuniculi* infection, increasing microbicidal activity, and reducing the fungal mechanisms involved in the evasion of the immune response.

**Highlights:**

- *E. cuniculi* phagocytosis and microbicidal activity by macrophages increases in the presence of B-1 cells
- M1 macrophage profiles were predominant in the presence of B-1 cells
- Extrusion of the polar tubule of *E. cuniculi* occur inside M2 macrophages in cultures without B-1 cells
- B-1 cells derived phagocytes (B-1CDP) identified with microbicidal activity against spores of *E. cuniculi*

**Author Summary:** The adaptive immune response plays a key role against *Encephalitozoon cuniculi*, an opportunistic fungus for T cells immunodeficient patients. The role of B cells and antibody play in natural resistance to *Encephalitozoon cuniculi* remains unresolved. Previously, we demonstrated that B-1 deficient mice (XID), an important component of innate immunity, were more susceptible to encephalitozoonosis, despite the increase in the number of CD4^+^ and CD8^+^ T lymphocytes. Here we observed that the absence of B-1 cells was associated with a larger population of M2 macrophages, an anti-inflammatory profile, which had lower microbicidal activity and phagocytic *E. cuniculi* spores were seen with the extrusion of the polar tubule, which is an important mechanism of evasion of the immune response. The results showed the importance of B-1 cells in the modulation of macrophage function against *E. cuniculi* infection, increasing microbicidal activity, and reducing the fungal mechanisms involved in the evasion of the immune response.

## Introduction

Microsporidia are obligate intracellular spore-forming microorganisms that can infect a wide range of vertebrate and invertebrate species. These fungi have been recognized as human pathogens and are particularly harmful to immunodeficient patients infected with HIV. Since then, interest among researchers of *in vitro* culture techniques has increased, with more people studying their biology and immune response against them [1].

*Encephalitozoon cuniculi* is one of the most common microsporidian species, in humans or animals. It is considered to be an emerging zoonotic and opportunistic pathogen in immunocompromised as well as immunocompetent individuals [2]. Spores of *E. cuniculi* can survive in macrophages, spread throughout the host, and cause lesions in organs of the urinary, digestive, respiratory, and nervous systems [3].

The adaptive immune response is critical for the elimination of *E. cuniculi*, but the innate immune response forms the first line of defense against these pathogens. The infection by *E. cuniculi* induces CD8^+^ cytotoxic T lymphocyte (CTL) response, which lyses the infected cells by perforin-dependent mechanisms [1]. Although antibody response during *E. cuniculi* infection has been recorded, it is clearly not sufficient to prevent mortality or cure the infection, making cell-mediated immunity critical for the survival of host infected by *E. cuniculi* [4]. The survival and replication of certain species of microsporidia within macrophages may be associated with the absence of phagosome-lysosome fusion [5]. Internalized microsporidium spores are normally destroyed within macrophages by the toxic activity of reactive oxygen and nitrogen species produced by the respiratory burst, and cytokines released by macrophages may be important in the protection against microsporidia [6].

B-1 cells are a subtype of B cells that account for 35%-70% of the B cells in the peritoneal cavity of mice [7]. They differ from B-2 cells in the expression of surface markers and function [8]. B-1 cells act as antigen-presenting cells, phagocytes, expressing myeloid (CD11b) and lymphoid markers (CD45/B220, CD5, CD19 and IgM), but not CD23, unlike B-2 cells [9]. The main function of B-1 cells in the innate immune system is the spontaneous secretion of natural antibodies, thereby maintaining immunoglobulin levels in the body without any stimulus or immunization [10]. In addition, B-1 cells also spontaneously secrete IL-10, while GM-CSF and IL-3 are secreted after lipopolysaccharide stimulation [10]. B-1 cells also regulate acute and chronic inflammatory diseases through the production of several immunomodulatory molecules, such as interleukin-10 (IL-10), adenosine, granulocyte-macrophage colony-stimulating factor (GM-CSF), IL-13, and IL-35, in the presence or absence of stimulus [11].

The possible role of B-1 cells in the dynamics of the inflammatory process of various etiologies is unknown and researchers have demonstrated the role of these cells in the functional regulation of macrophages. Also, B-1 cells are able to differentiate into phagocytes (B-1CDP), characterized by the expression of F4/80 and increased phagocytic activity [12,13]. Furthermore, Popi *et al.* [14] demonstrated that BALB/c mice were more susceptible to *Paracoccidioides brasiliensis* infection compared to XID (B-1 cell deficient) mice and attributed the down-regulation of macrophage function to IL-10 secreted by B-1 cells.

In recent studies from our group, we demonstrated that B-1 cell deficient XID mice were more susceptible to experimental encephalitozoonosis than BALB/c mice, evidenced histologically with more prominent inflammatory lesions and fungal burden [15,16]. Although the mechanism of action of B-1 cells in the resistance of BALB/c mice to encephalitozoonosis is not fully understood, a significant increase in the population of peritoneal macrophages was reported in BALB/c mice infected with *E. cuniculi* [15]. We hypothesized that B-1 cells from the peritoneal cavity (PerC) could differentiate in infected animals into B-1 cell-derived phagocytes (B-1 CDP), which could then promote the phagocytosis of *E. cuniculi* spores and also influence the macrophage function in this context. Herein, we tested this hypothesis using the ultrastructure and phenotypic analysis of adherent peritoneal cells (APerC) to evaluate their *in vitro* behavior in BALB/c and B-1 cell-deficient XID mice against the fungus *E. cuniculi*. The presence of B-1 cells facilitated the phagocytic and microbicidal activity and increased apoptosis in the APerC cultures. These cells were associated with the presence of M-1 macrophages with increased proinflammatory cytokine production. Electron micrographs showed an intimate physical relationship between B-1 cells and macrophages, indicating communication and modulation of activity, demonstrating that the presence of B-1 cells drives the behavior of macrophages and consequently the innate immune response in encephalitozoonosis.

## Results

### B-1 cells increase macrophage activity in *E. cuniculi* infection

To assess the influence of B-1 cells on the activity of macrophages in *E. cuniculi* infection, APerC obtained from PerC of B-1 cell-containing BALB/c and B-1 cell-deficient XID mice were co-infected with *E. cuniculi* for ultrastructural analyses at 1, 48, and 144 h. The phagocytosis of spores occurred in cell cultures of both BALB/c and XID APerC groups, via actin rearrangement characterized by projections of the cellular membrane of macrophages near or around the spores (Figures 1A and 1B), and remained within phagosome vacuoles that were dispersed throughout the cytoplasm. XID macrophages showed large numbers of vacuoles in the cytoplasm and several membrane projections (pseudopods), indicating widespread and strong phagocytic activity (Figure 1A).

**Figure 1.**
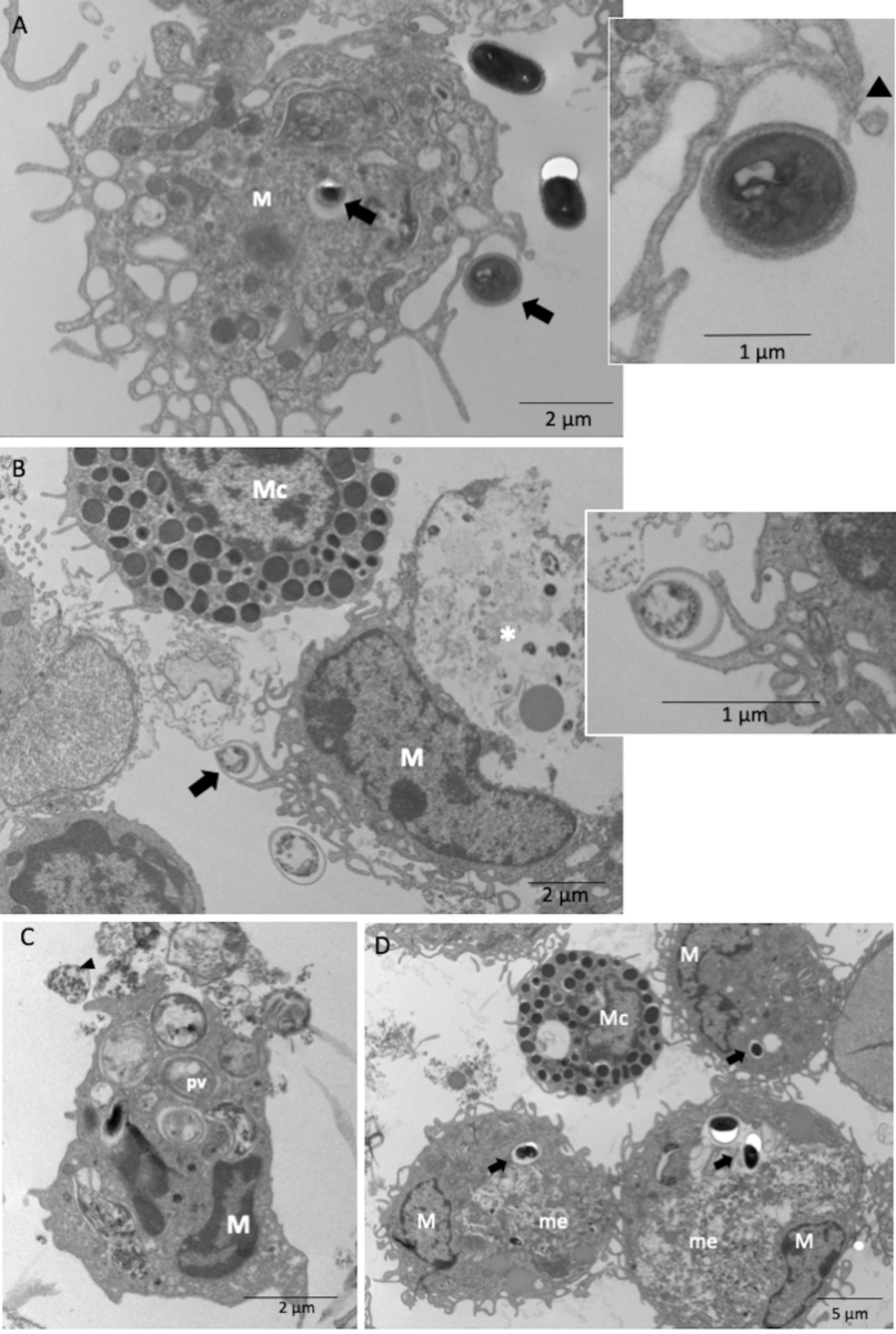
Microbicidal activity of BALB/c and XID APerC after 1 h of infection with *E. cuniculi* spores by ultramicrography. (A and insert) Projections of the cellular membrane (head arrows) of macrophages near or involving the spores (arrow) of *E. cuniculi* in XID APerC. (B and insert) Projections of the cellular membrane of macrophages involving *E. cuniculi* spore (arrow) of in BALB/c APerC and amorphous material (*) inside phagosome vacuole. Note the presence of mast cell. (C) Homotypic phagosomes vacuoles (pv) with *E. cuniculi* spores inside and amorphous material exocytosed (head arrow) in BALB/c APerC macrophages. (D) Homotypic phagosomes vacuoles with intact spores (arrow) and megasome (me) with amorphous material inside in the macrophages (M) from XID APerC. Note mast cells (Mc) in contact with macrophages.

Intact and degenerating internalized spores surrounded by a vacuolar membrane, which is a typical phagosome vacuole, were observed (Figures 1C). Under normal circumstances, the interaction of phagosomes with each other and with other organelles is tightly regulated, and it is well documented that phagosomes containing inert particles are not subjected to homotypic fusion [17]. The important finding of this study is the formation of megasomes by the fusion of homotypic phagosomes containing a single *E. cuniculi* spore (Figure 1D). Therefore, the ability of *E. cuniculi* to induce the fusion of phagosomes resembles that of *Helicobacter pylori* [18] and *Chlamydia trachomatis* [19].

In AperC cultures of both BALB/c and XID mice, the microbicidal activity was identified by the presence of a large amount of amorphous and electron-dense material either inside the phagocytic vacuoles and megasome in the cytoplasm of macrophages or undergoing exocytosis (Figures 1B and 1D). The same findings were seen at 48 h (Figure 2A–2D) with an increase in the ratio of degenerated spores, and at 144 h with only a few intact spores inside the phagocytic cells in case of XID APerC (Figure 2E and 2F). Myelin figures indicating spore degeneration were recorded (Figure 2B). Another important finding was the presence of degenerating macrophages (Figure 2D). At 144 h, the APerC of BALB/c showed an absence of macrophages and mature spores of *E. cuniculi* outside the cells (Figures 2G and 2H) and degenerated lymphocytes. The intracellular proliferative stages of *E. cuniculi* (meronts, sporonts, or sporoblast) were not observed.

**Figure 2.**
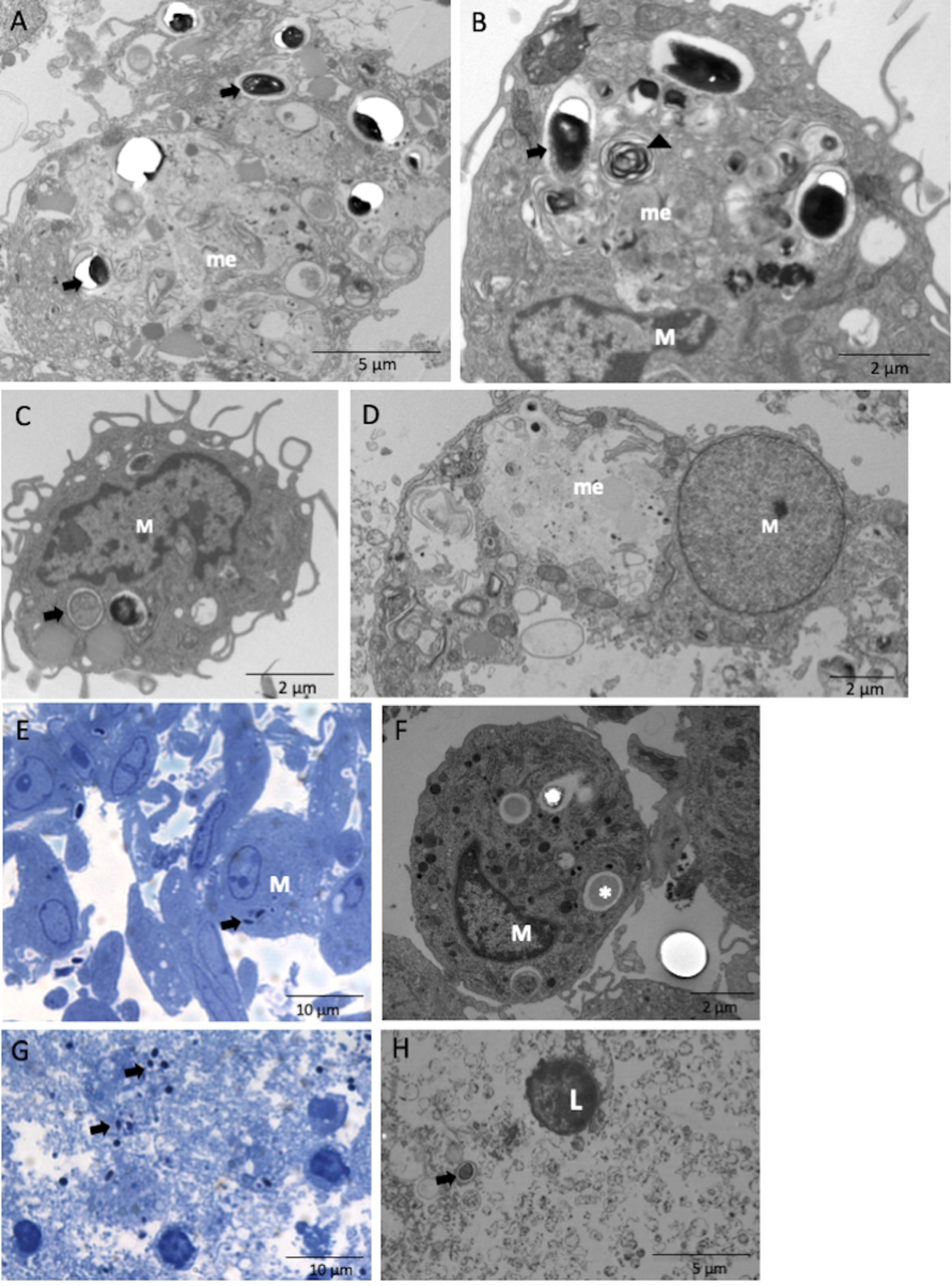
Microbicidal activity of BALB/c and XID APerC after 48 and 144 h of infection with *E. cuniculi* spores. (A) Megasome (me) with amorphous and electron dense material and *E. cuniculi* spores (arrow) inside. Ultramicrography of macrophages from APerC BALB/c after 48 h of infection. (B) Megasome with amorphous and electrodense material, myelin figure (head arrow) and spores (arrow) in XID APerC by ultramicrography after 48 h of infection. (C) Homotypic phagosomes vacuoles in macrophages from APerC BALB/c after 48 h of infection, by ultramicrography. (D) Degraded macrophage (M) containing amorphous material and involving spore of *E. cuniculi* in megasome (me), by ultramicrography. (E) Photomicrography of phagocytic cells with *E. cuniculi* spores inside them (arrow) in APerC XID with144 h of infection. (F) Ultramicrography of a phagocytic cell with *E. cuniculi* spores in lysis (arrow) in APerC XID after 144 h of infection. (G) Photomicrography of APerC BALB/c in the absence of macrophages and mature spores of *E. cuniculi* outside the cells (arrow) and degenerated lymphocytes (head arrow) after 144 h of infection. (H) Ultramicrography of APerC BALB/c with the absence of macrophages and mature spores of *E. cuniculi* outside the cells (arrow) and degenerated lymphocytes (head arrow) with pyknotic nucleus after 144 h of infection.

Using Calcofluor stain, we observed increased phagocytic capacity and index in BALB/c APerC compared to XID APerC (Figure 3), suggesting the involvement of B-1 cells in the increase in phagocytic activity. After 1 h, large numbers of preserved spores were observed within the macrophages from BALB/c and XID, but after 48 h, there were fewer intact spores within the phagocytic vacuoles from BALB/c APerC, while the spores remained preserved in XID APerC.

**Figure 3.**
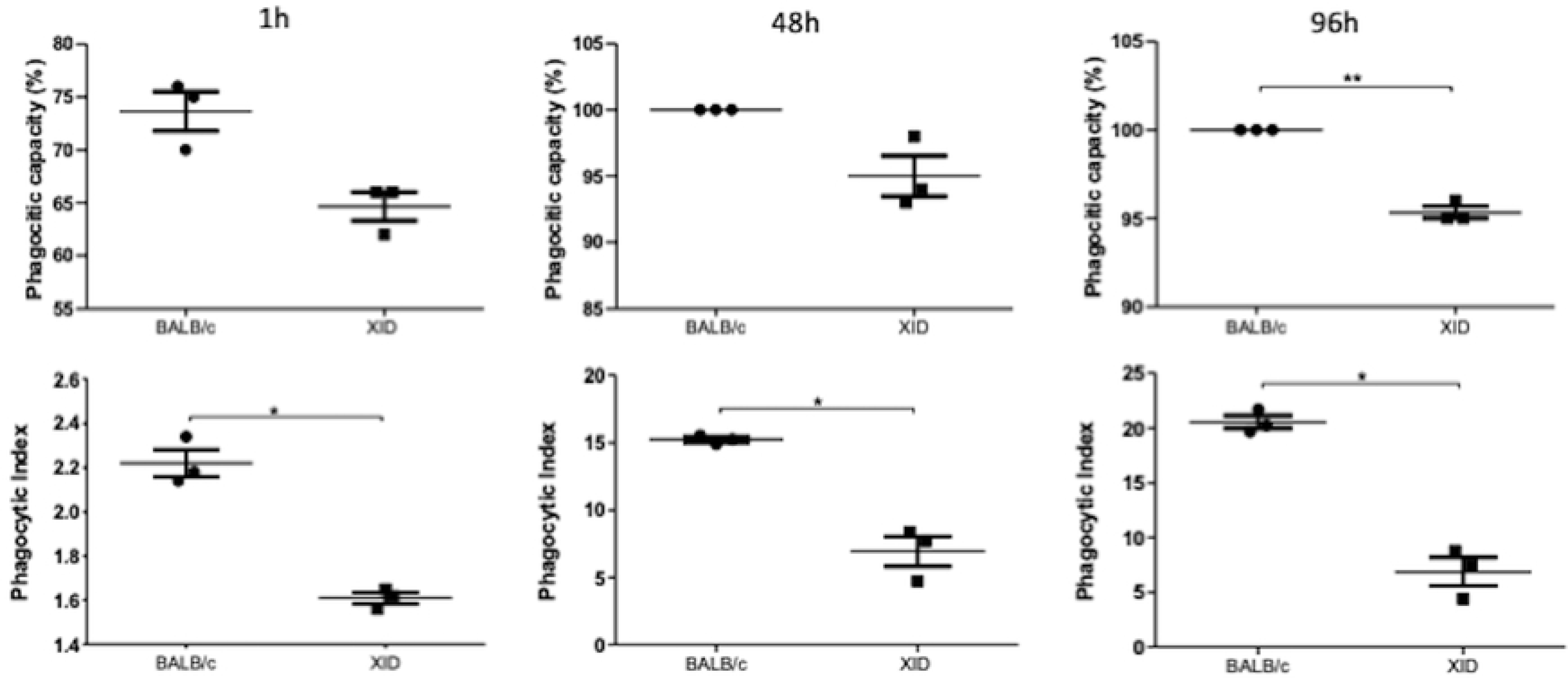
Phagocytic activity of BALB/c and XID mice adherent peritoneal cells infected with *E. cuniculi*. The phagocytic capacity and phagocytic index were obtained from adherent peritoneal cells infected with *E. cuniculi* after 1, 48, and 96 h. Date are represented as mean ± SEM (*p < 0.05, **p < 0.01, T test)

### Presence of B-1 cells reduces death by apoptosis in *E. cuniculi* infected APerC

As the ultrastructural analysis revealed that the BALB/c APerC phagocytes were no longer viable after 144 h, we evaluated the level of apoptosis and necrosis after 48 and 96 h for all experimental groups in order to examine the possible influence of *E. cuniculi* in the process of cell death (Figure 4). Given the abundance of cell death accompanying intracellular infection, two possibilities have been described as evasion mechanisms: 1) the pathogen can be destroyed along with the engulfed apoptotic cell (host antimicrobial activity) or 2) the pathogen can use the process of efferocytosis to disperse into new cellular hosts (“Trojan Horse” model) [20]. About 50–60% of cell death was observed in BALB/c AperC, which was higher than that in XID APerC (30–40%) after 48 h (Figure 4A). At 96 h, the percentage of the death of uninfected BALB/c and XID APerC was similar (about 70%), but a 40 to 50% reduction in the rate of cell death in the infected groups was evident (Figure 4A). In addition, after the infection, the BALB/c and XID APerC behaviors were antagonistic and the rate of apoptosis identified in the infected BALB/c APerC was higher than that observed in XID APerC at both the time intervals (Figure 4B). Similar to our findings, Martin *et al.* [20] reported that *M. tuberculosis*-infected macrophages that die by apoptosis are rapidly engulfed by uninfected macrophages via efferocytosis and this process is responsible for the bactericidal effect associated with apoptosis.

**Figure 4.**
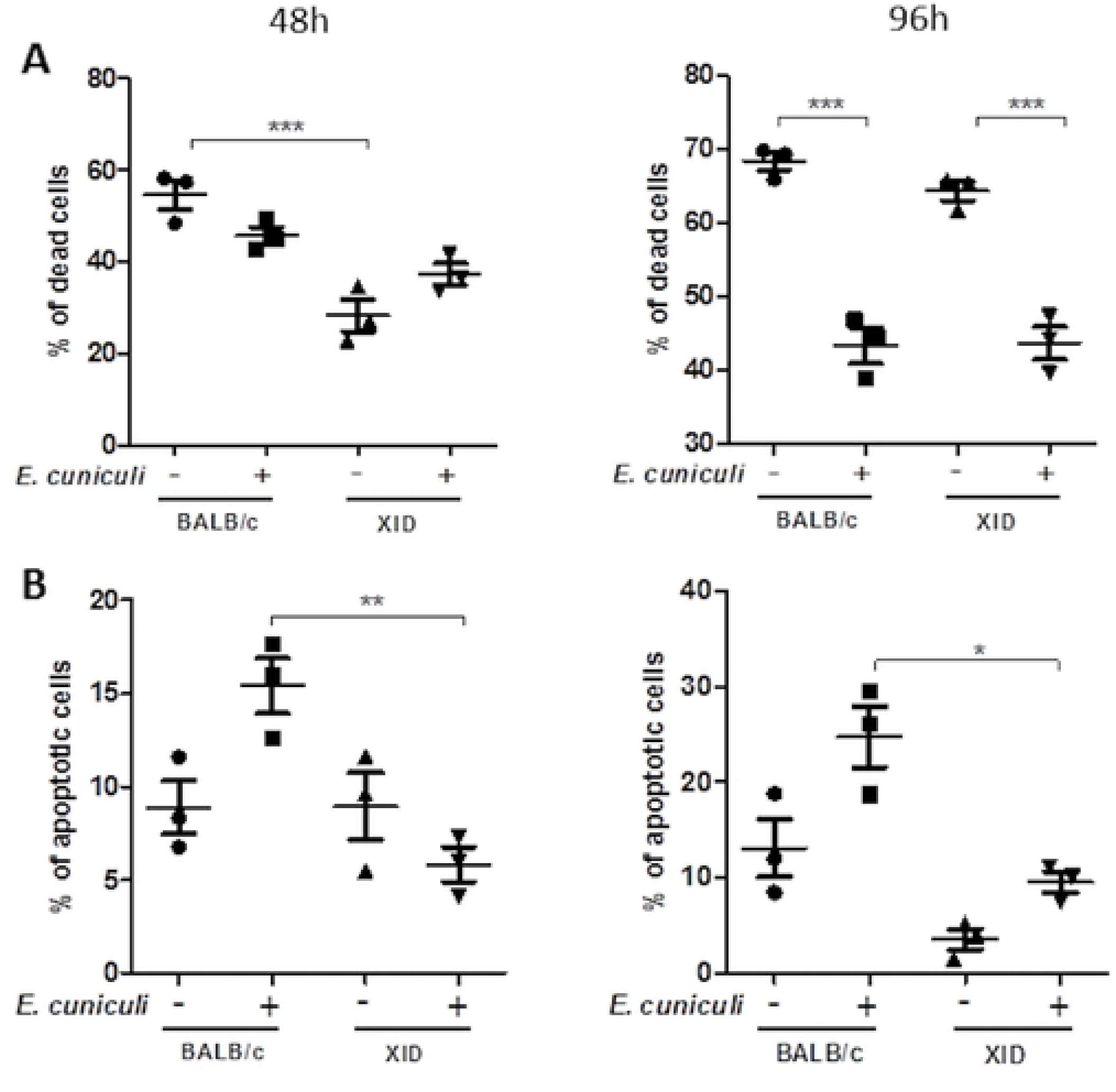
Necrosis and apoptosis of BALB/c and APerC cultures at 48 and 96 h. (A) % of death cells. (B) % of apoptotic cells. The data are presented as mean ± SEM (*p < 0.05, **p < 0.01, t-test)

The spores that were internalized by macrophages in XID APerC cultures had a contact area between the phagocytic vacuolar membrane of the cell and the spore wall (Figures 5A and insert), suggesting intimate contact with the phagocytic cells. Furthermore, we observed the extrusion of the polar filament by an internalized spore from the phagosome vacuole at 1 h (Figures 5B and insert). The ejection of the polar filament from the phagosome into the cell has been reported in the literature [1], but has been rarely demonstrated by ultrastructure analysis. This finding suggests an evasion of the microbicidal activity. Therefore, we hypothesize that the absence of B-1 cells could direct the macrophages to M2 profile and favor the dissemination of *E. cuniculi*.

**Figure 5.**
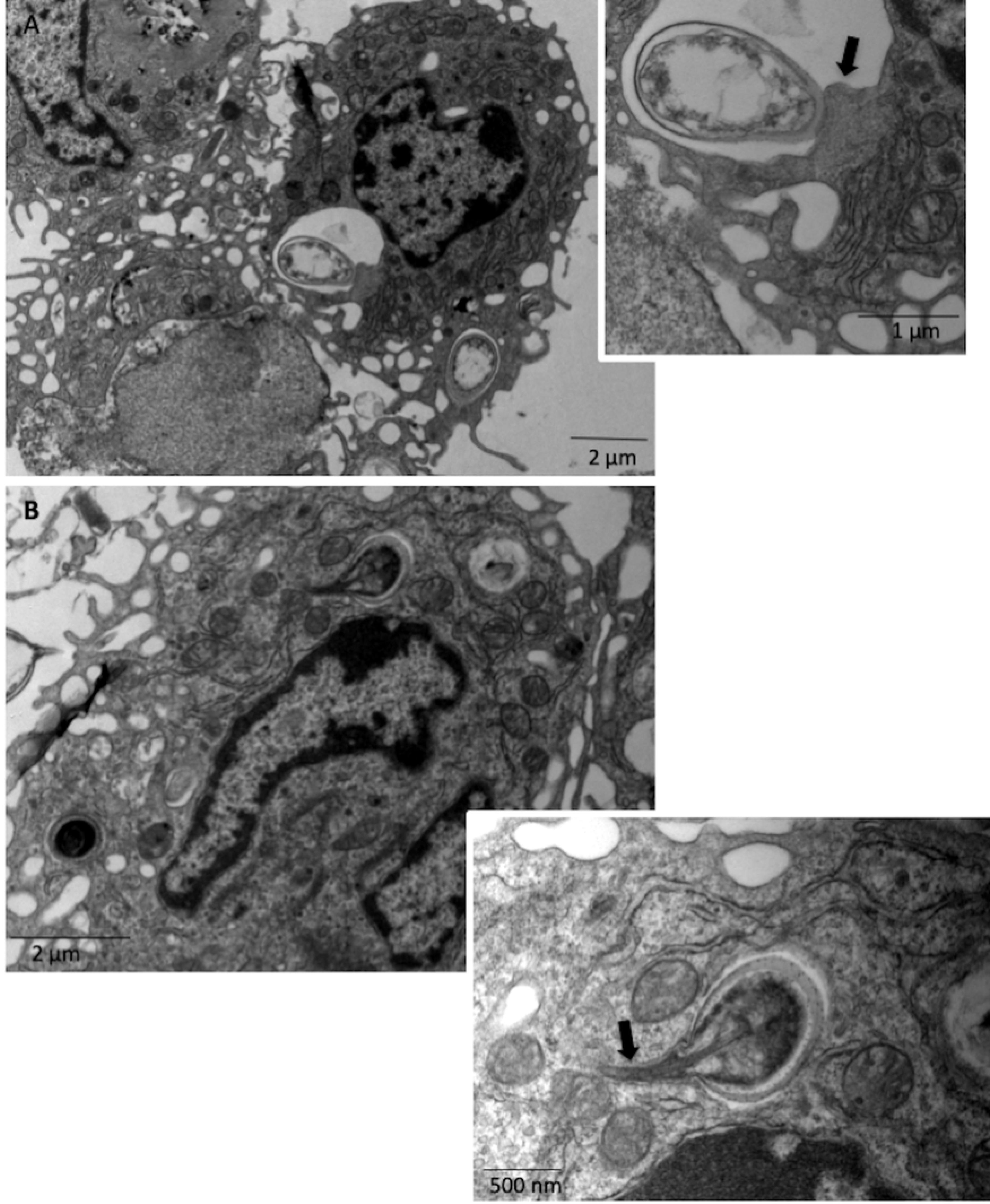
Relations between phagocytes from XID APerC and phagocytized *E. cuniculi* spores after 1 h of infection by ultramicrography. (A–B) Contact area between the phagocytic vacuole membrane of the macrophages and the internalized spore wall (arrow). (C–D) Germinated spore with extruded polar tube (arrow) from the phagosome vacuole.

To test this hypothesis, we evaluated the expression of CD40 and CD206 in macrophages from BALB/c and XID APerC cultures, which may suggest the classical activation-M1 profile (CD40^+high^ CD206^+low^) or the alternate activation-M2 profile (CD206^+high^ CD40^+low^) process. We also analyzed the mean fluorescence intensity of CD80 and CD86 co-stimulatory molecules to measure the activation of these macrophages and aid in the characterization of M1/M2.

### B-1 cells are associated with M1 macrophage profile

The proportion of macrophages in XID APerC (approximately 60%) was significantly higher than that in BALB/c APerC (approximately 20%), regardless of microsporidial infection (Figure 6A). Since the BALB/c culture also contains B-1 cells that adhere together with the macrophages in the first incubation [15,16], this result was expected. After infection, the percentage of macrophages in XID APerC was 80%, which is higher than that observed in uninfected cultures, both at 48 and 96 h (Figure 6A). In BALB/c APerC, no difference was observed in this parameter. At 48 h, approximately 80% of the BALB/c APerC macrophages expressed CD40^+^ while approximately 40% of XID APerC macrophages expressed CD40^+^ (Figure 6B). At 96 h, the percentage of these cells decreased to about 60% of CD40^+^ cells in infected BALB/c AperC and about 40% in infected XID APerC. At 48 h, the CD206^+^ expression on macrophages was higher in XID APerC than in BALB/c APerC (Figure 6C). At 96 h, there was no difference between the groups. We also analyzed the ratio of CD40^+^ cells to CD206^+^ cells and observed a higher ratio in the infected BALB/c than in XID (Figure 6D). In addition, we observed that infected BALB/c APerC macrophages expressed more CD80 and CD86 costimulatory molecules (Figure 6E) than macrophages from the other groups. These molecules are expressed on the surface of activated macrophages, especially M1 macrophages, which present a higher level of expression. These results together indicate a higher proportion of M1 profile macrophages in BALB/c APerC and M2 profile in XID AperC. The prevalence of *E. cuniculi* infection reinforces these profiles.

**Figure 6.**
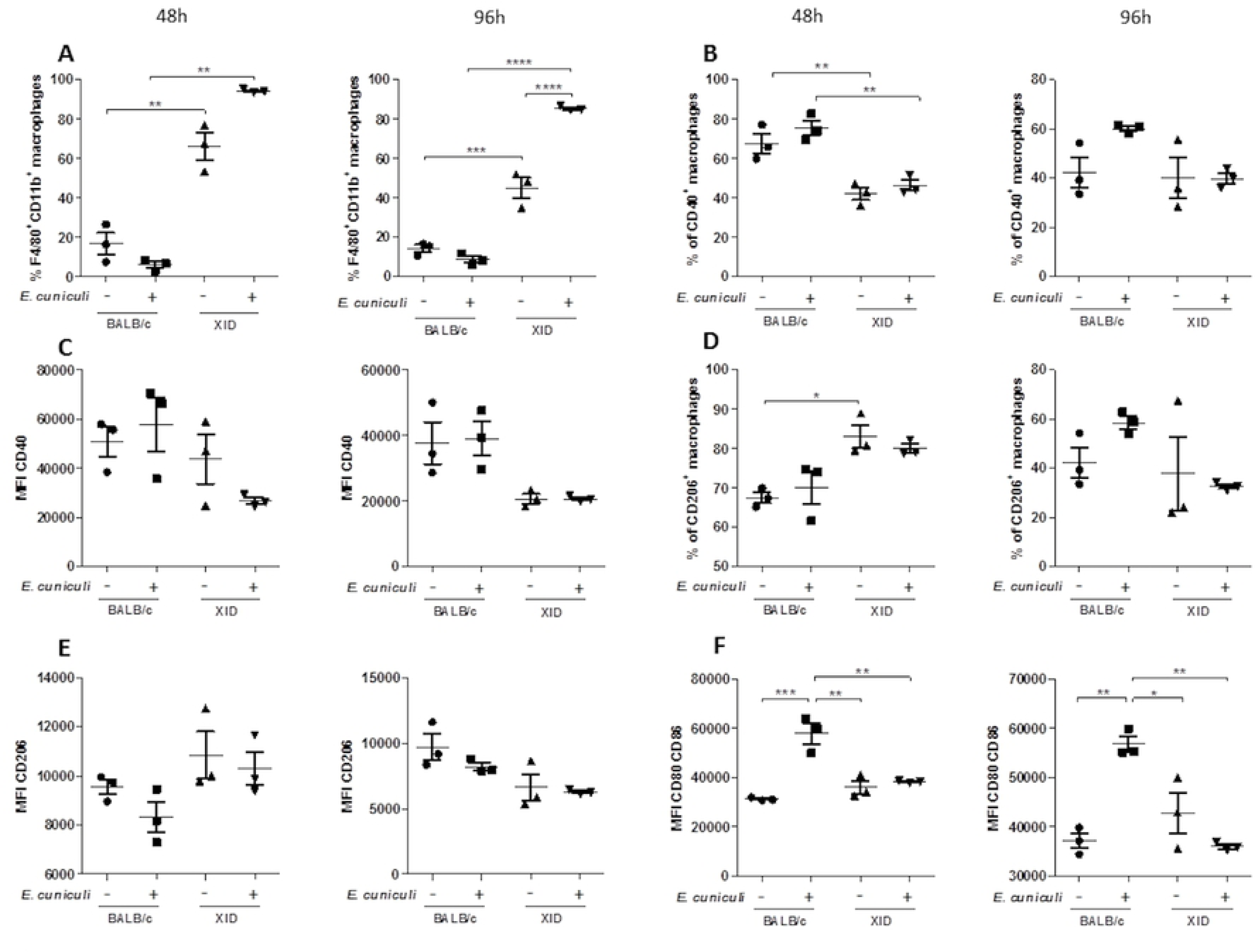
Activation phenotype of macrophages from BALB/c or XID APerC infected with *E. cuniculi* spores. (A) Total percentage of F4/80^+^ and CB11^b+^ macrophages for each group. (B) Proportion of F4/80^+^ and CB11^b+^ macrophages expressing CD40^+^. (C) Mean fluorescence intensity (MFI) of CD40 molecule on macrophage. (D) Proportion of F4/80^+^ and CB11^b+^ macrophages expressing CD206. (E) Mean fluorescence intensity (MFI) of CD206 molecule on macrophage. (F) Mean fluorescence intensity of CD80^+^ and CD86^+^ on macrophages. The date are represented as mean ±SEM (* p <0.05, ** p <0.01, *** p <0.001, **** p <0.0001, Two-way ANOVA with multiple comparisons and Bonferroni post-test).

### B-1 cells reduce NO production by macrophages but increase proinflammatory cytokines in ***E.*** *cuniculi* infection

The levels of NO in BALB/c APerC infected with *E. cuniculi* were similar to those of the uninfected group at all intervals. On the other hand, the level of NO in infected XID APerC was significantly higher than that in the uninfected group at 96 h (Figure 7). These findings suggest that the presence of B-1 cells has an inhibitory effect on NO production by macrophages, which indicates that the elimination of *E. cuniculi* by macrophages involves another mechanism. Also, while the XID APerC contained predominantly M2 macrophages, the NO production increased after *E. cuniculi* infection.

**Figure 7.**
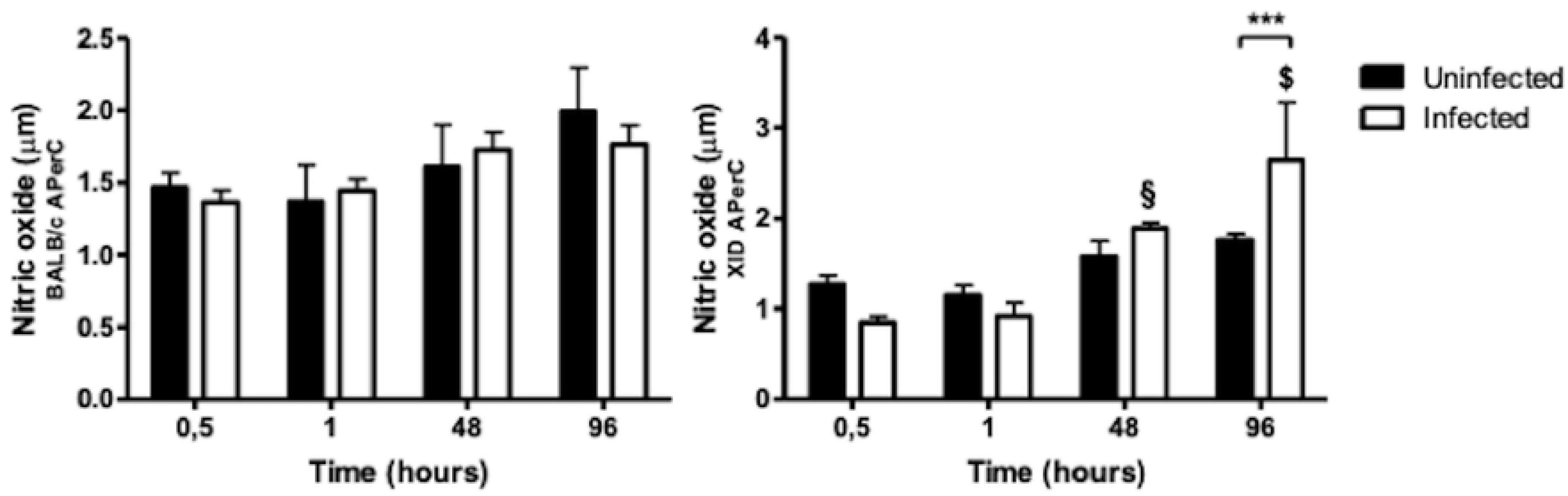
Levels of nitric oxide (NO) in supernatants of BALB/c and XID APerC. Date are presented as mean ±SEM (# p<0.05 compared to 1 h and 0.5 h, $–p<0.05 compared to other times, §–p<0.01 compared to other times, one-way ANOVA test with the Bonferri post-tests shows)

BALB/c APerC demonstrated a proinflammatory cytokine profile characterized by increased IL-6 and TNF-α at 96 h of infection, although a reduction of IL-6 was observed in the previous time intervals (Figure 8). In contrast, we observed a reduction in the proinflammatory cytokines (IL-6 and TNF-α) and anti-inflammatory IL-10 at 96 h of infection in the XID APerC supernatant (Figure 8). Although the cytokine IL-12 was tested in this study, it could not be detected. The levels of IFN-γ were not statistically different between the tested groups in across the time intervals (data not shown). These findings are in line with the observed macrophage profiles, with a predominance of M1 in BALB/c APerC and production of proinflammatory cytokines. The levels of MCP-1 decreased in BALB/c APerC across the time intervals observed, but interestingly, the XID APerC did not appear to produce this cytokine at 96 h. This result could suggest the recruitment of new macrophages to the environment in *in vivo* conditions.

**Figure 8.**
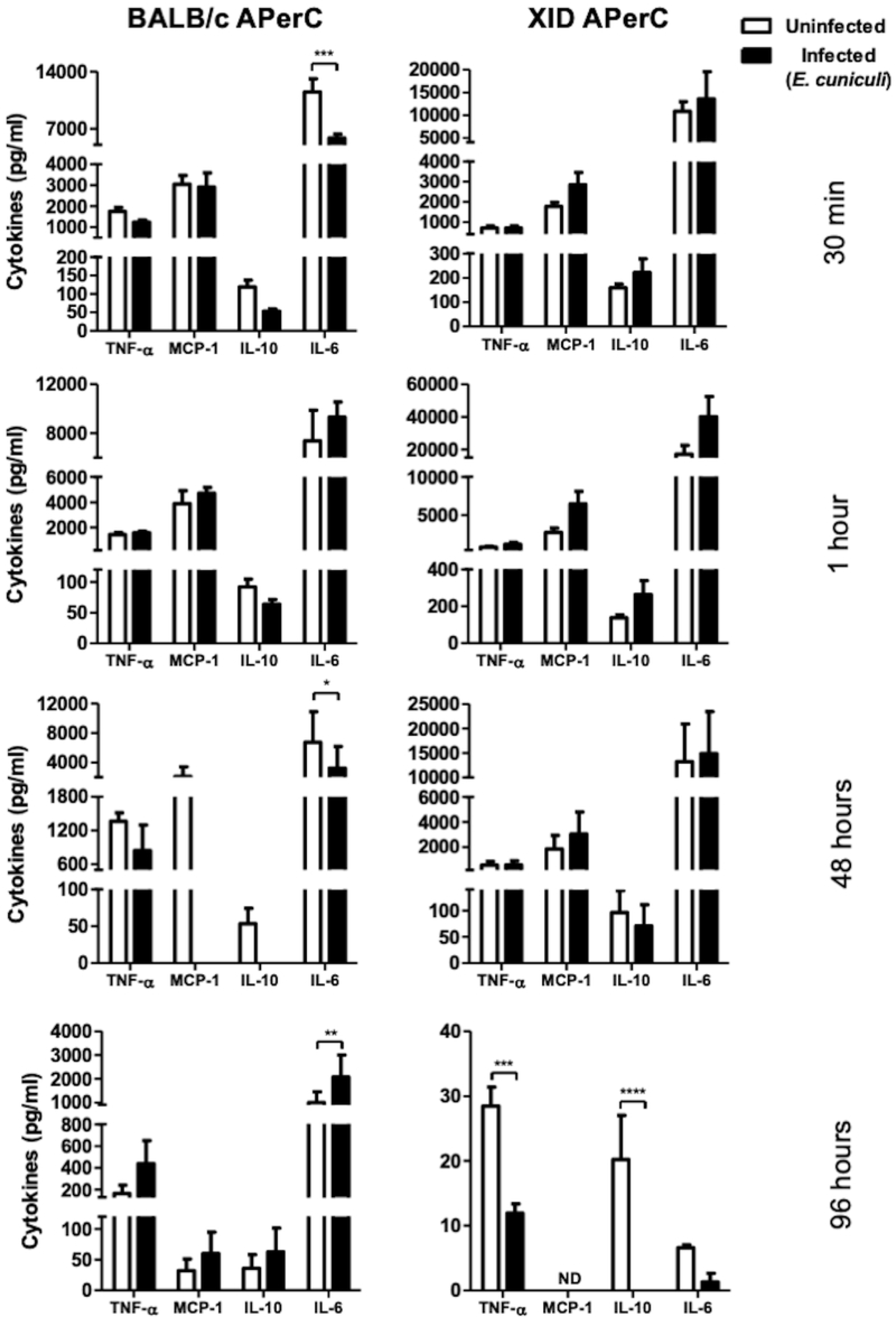
Cytokine levels in the supernatants of BALB/c and XID APerC. Date are represented as mean ±SEM (* p <0.05, ** p <0.01, *** p <0.001, **** p <0.0001 and ND = not detected, Two-way ANOVA with multiple comparisons and the Bonferroni post-test).

### Contact and Interaction Between Peritoneal Cells

The B-1 cells were identified in BALB/c APerC by their ultrastructural morphological characteristics such as a lobed nucleus with bridges of the nuclear membrane joining the lobules and a well-developed endoplasmic reticulum with few mitochondria [21] (Figure 9A and 9B). We also observed the presence of cytoplasmic projections of pre-B-1 CDP cells and B-1 lymphocytes adhered to or near extracellular *E. cuniculi* spores in BALB/c APerC (Figure 9B). Intercellular communication involving other lymphocytes (Figure 9A) and mast cells in BALB/c (Figure 9C) and XID APerC (Figure 1D), and plasma cells (Figure 9D) in BALB/c APerC were often observed.

**Figure 9.**
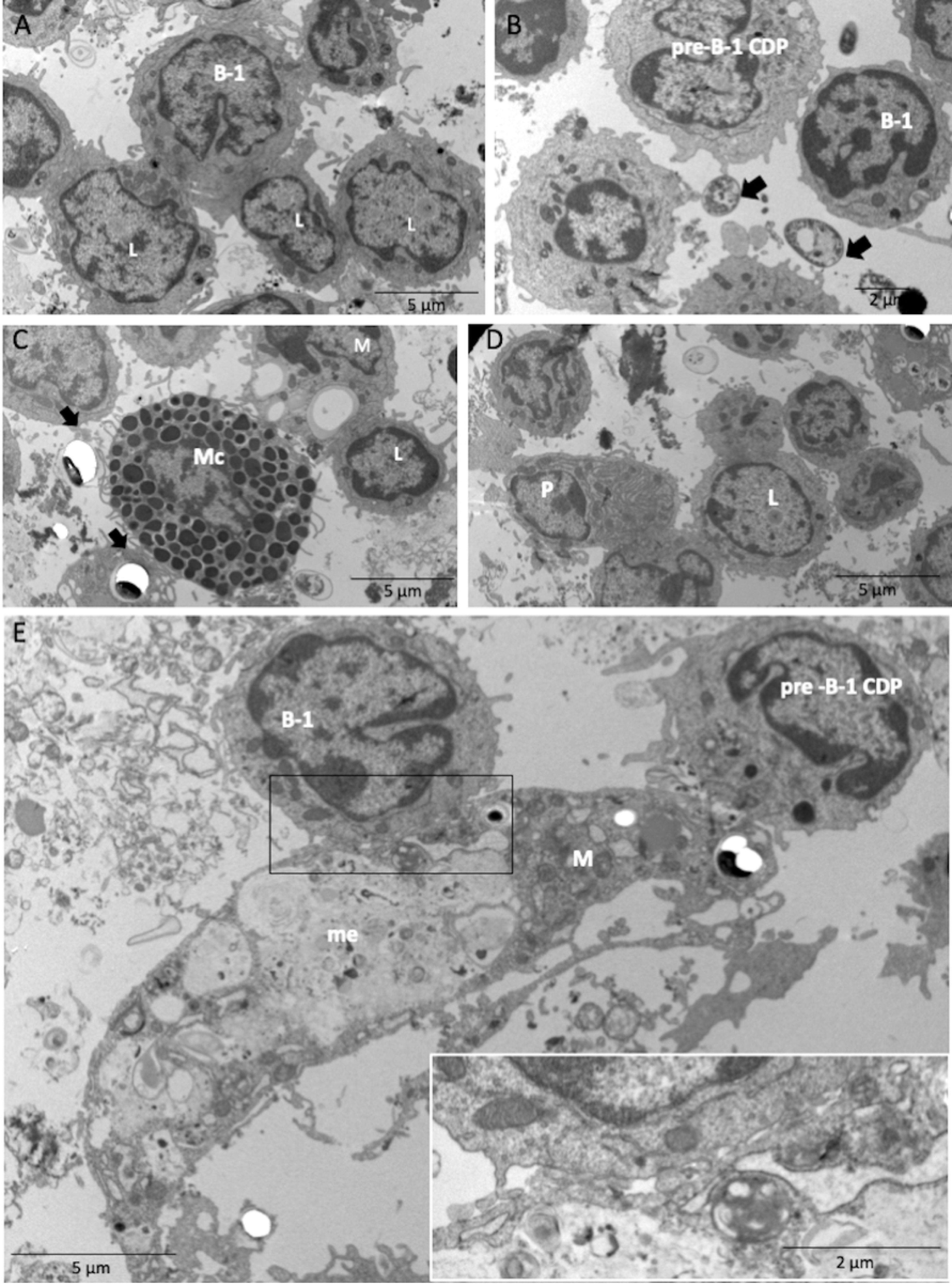
Cross-talk between the peritoneal cells of BALB/c AperC after infection with *E. cuniculi* spores by ultramicrography. (A) Typical B-1 cell (B-1) with a lobular organization of the nucleus in communication with others lymphocytes (L). (B) Pre-B-1 CDP in contact with B-1 cells and *E. cuniculi* spores (arrow). (C) Mast cell (Mc) contact macrophages (M) and lymphocytes (L). (D) Plasma cell (P) in communication with lymphocytes (L). (E) B-1 cells (B-1) and pre-B-1 CDP in communication with macrophage with megasome (me). Insert: Note the close relationship between macrophage membranes and B-1 cell.

### The activity of B-1 Cell-Derived Phagocyte in *E. cuniculi* infection

It was previously demonstrated that B-1 cells differentiate to acquire a mononuclear phagocyte phenotype following attachment to a substrate *in vitro*, which are named B-1 cell-derived phagocytes (B-1CDPs) [22,23]. It has already been demonstrated that these cells are able to phagocytose various pathogens *in vivo* and *in vitro* [23,24,25,26]. We used B-1 CDP cultures to evaluate the participation of B-1-derived phagocytes in encephalitozoonosis. In B-1 CDP cultures, we observed B-1 CDPs and B-1 cells with preserved characteristics after 1 h and 48 h (Figure 10), indicating that a part of the B-1 cells had become phagocytes. Mussalem *et al.* [27] demonstrated that infection with *Propionibacterium acnes* induced the commitment of B-1 cells to the myeloid lineage and their differentiation into phagocytes. We observed the contact of spores with B-1 cells and pre-B-1-CDPs (Figure 10A). B-1 CDPs with spores of *E. cuniculi* in the process of lysis within phagocytic vacuoles, amorphous material in megasomes (Figure 10B), and intact spores were also visualized in the B-1 CDP cultures (Figure 10C). Other authors also demonstrated the phagocytic and microbicidal ability of peritoneal B-1 cells [28,29]. The ultrastructure of *E. cuniculi* spores was typical of non-germinated mature spores and showed of a thick wall composed of an electron-dense outer layer (exospore), an electron-lucent inner layer (endospore), and a plasma membrane enclosing the cytoplasm (Figure 10C). B-1 cells and pre-B-1 CDP have abundant microvesicles in their membranes (Figures 10D and 10E), indicating exocytosis. Exosomes are nanosized membrane microvesicles that have the capability to communicate intercellularly and to transport cell components [30].

**Figure 10.**
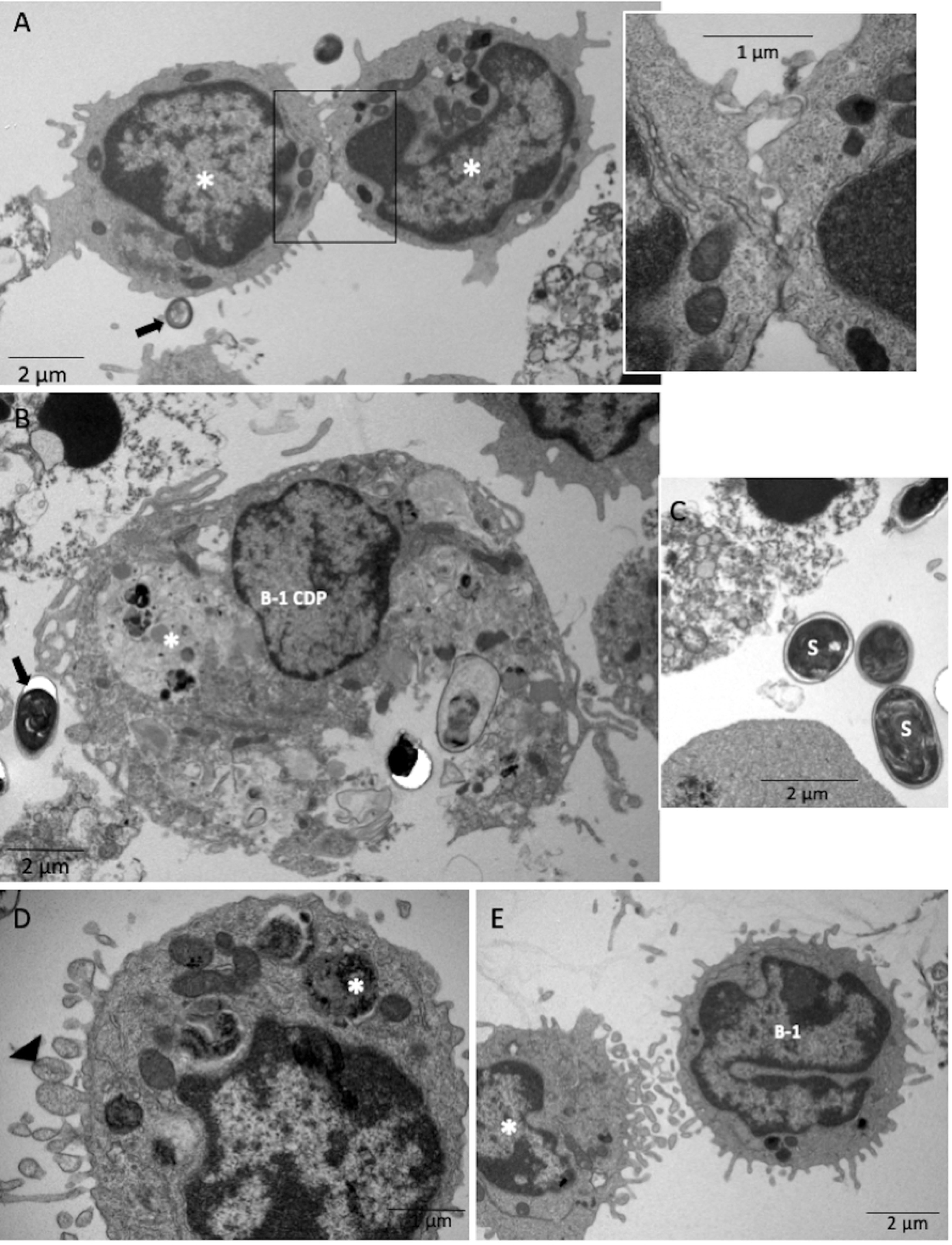
Phagocytic activity of B-1 CDP culture after 1 h and 48 h of infection with *E. cuniculi* by ultramicrography. (A) Pre-B-1 CDP (*) in communication between them and with *E. cuniculi* spores (arrow). Amplified view of the interaction between B-1 cells. (B) B-1 CDP with the presence of amorphous material inside phagocytic vacuoles (*). (C) Non-germinated spores (s) outside the cells with a thick wall composed of two layers and a plasma membrane. (D) Microvesicles (head arrow) in the membrane of pre-B-1 CDP with the presence of amorphous material (*) inside. (E) Microvesicles in membranes of pre-B-1 CDP (*) and B-1 cell.

Thus, here we identified two forms of intercellular communication through contact and release of extracellular vesicles (Figures 10A and 10E, respectively). In B-1 CDP cultures, infection was associated with a reduction in the percentage of cell death at 96 h and death by apoptosis at 48 h (Figure 11A). Spores outside the cells were identified under light microscopy (data not shown) and low index of phagocytosis was determined after one hour with an increasing trend across the time intervals (Figure 11B). The levels of NO in the supernatants of B-1 CDP cultures infected with *E. cuniculi* were not different from that of the uninfected group at all the time intervals (Figure 11C). We observed a considerable increase in the proinflammatory cytokine TNF-α at 30 min and 1 h in the infected cultures, while the TNF-α and MCP-1 cytokine levels decreased at 96 h (Figure 11D). Cytokine IL-12 was not detected. IFN-γ levels did not statistically vary across the time intervals (data not shown).

**Figure 11.**
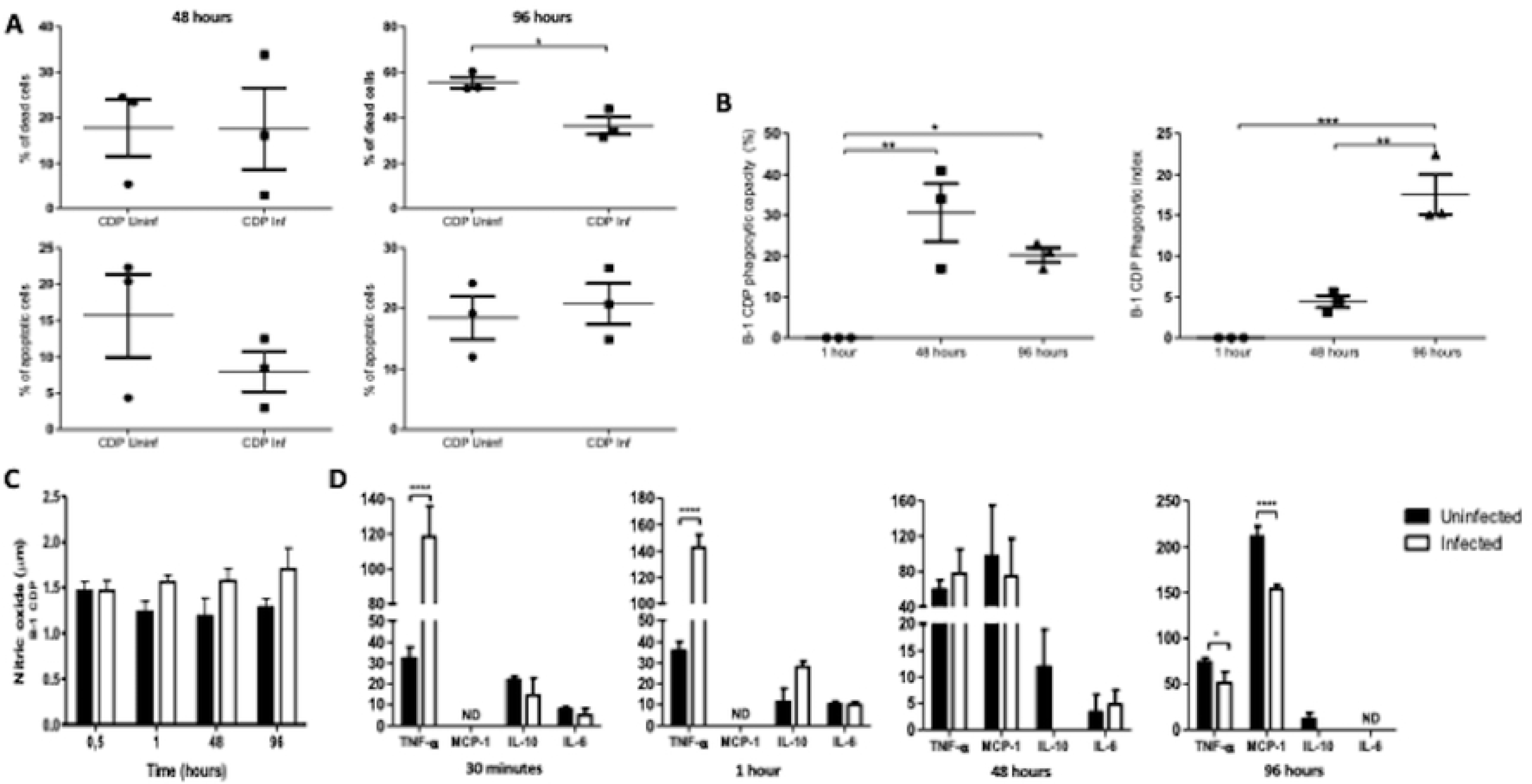
NO and cytokine levels in the supernatants from B-1 CDP cultures. Phagocytic index (A) NO levels. Two-way ANOVA test with Tukey’s post-test. (B) Cytokines levels. Two-way ANOVA test with multiple comparisons and the Bonferroni post-tests at *p<0.05, ****p<0.0001 and ND = not detected.

## Discussion

The role of B-1 cells in immunity against fungal, protozoan, bacterial, and helminthic infections has been described by us as well as by other researchers. Popi et al. [14] demonstrated that BALB/c mice were more susceptible to experimental infection by *Paracoccidioides brasiliensis* than XID mice, suggesting that B-1 cells may favor infection. This was consistent with other experimentally induced infections, such as *Mycobacterium bovis* bacillus Calmette-Guerin (BCG) [31] and the *Trypanosoma cruzi* [32]. B-1 cells secrete IL-10 and use it as an autocrine growth factor. Popi et al. [12] demonstrated *in vitro* that this cytokine decreases the production of nitric oxide and hydrogen peroxide by macrophages, which decreases their phagocytic capacity, compromising the innate immune response and antigenic presentation.

Nevertheless, B-1 cells are critical for the early control of infections with encapsulated bacteria such as *Streptococcus pneumoniae* [33], viruses such as the *Influenza* virus [34], or fungi [35,36]. In line with these studies, our group demonstrated that XID mice were more susceptible to encephalitozoonosis than BALB/c mice, suggesting that the pathogenicity caused by *E. cuniculi* depends on the relationship between the multiplication of parasites and the host immune response [15,16]. In addition, da Costa *et al.* [15] showed that BALB/c mice infected with *E. cuniculi* showed an increase in the number of macrophages and plasma levels of IFN-γ, which was responsible for the activation of macrophages and elimination of the pathogens. Confirming this, here we demonstrated *in vitro* that B-1 cells upregulated the macrophage activity against *E. cuniculi*, characterized by higher phagocytic index and microbicidal capacity, as well as increased macrophage death. The presence of intimate contact between B-1 cells and macrophages suggested communication between these cells and modulation of activity. In addition, B-1 CDP also showed intense phagocytic and microbicidal activities, a fact that may explain the numerical increase of macrophages in BALB/c mice, as previously described by our group [15].

The plasticity of macrophages has been demonstrated over the last few years. Depending on the stimulus received by macrophages (pathogens or injured tissue), their receptors trigger a decision to kill (M1) or repair (M2) [37]. The data from transcriptome analysis demonstrated the existence of distinct polarization phenotypes of macrophages associated with specific pathological conditions [38,39]. It is widely accepted that the phenotype of macrophages reflects the immediate microenvironment, wherein other cells can participate in this process. In addition, the role of B-1 cells in the polarization of peritoneal macrophages to an M2 profile in tumor condition was proposed by Wong et al. [40]. In contrast, we demonstrated that macrophages from BALB/c APerC present M1 profile (CD40 ^high^ CD206 ^low^CD80/86 ^high^) while macrophages from XID APerC present M2 profile (CD206 ^high^ CD40^low^CD80/86^low^). These results may partly explain the susceptibility of XID mice to encephalitozoonosis when compared to BALB/c mice, as previously demonstrated by our group [15]. As demonstrated in this study, M1 macrophages possess intense phagocytic activity and capacity to produce proinflammatory cytokines. The presence of megasomes, amorphous material, degenerated spores, and myelin figures within phagocytes in BALB/c APerC confirmed the high microbicidal potential of these macrophages. Furthermore, the expression of CD80 and CD86 produces a second signal necessary for the proliferation and activation of T lymphocytes, a fact that demonstrates the importance of innate response in the production of acquired effective response against microsporidia. AperC infected with *E. cuniculi* showed higher expression of these molecules.

M2 macrophages produce ornithine that promotes proliferation and tissue repair, increasing levels of TGF-β, IL-10, IFN, chitinases, matrix metalloproteinases, scavenger receptors, and have a poor microbicidal function [37,41]. Herein, apart from the M2 profile phenotypically associated with XID APerC macrophages, we observed a lower rate of phagocytosis and delayed microbicidal activity associated with reduced pro-inflammatory cytokines, a phenomenon linked with the absence of B-1 cells.

The infection process of *E. cuniculi* involves the forced eversion of a coiled hollow polar filament that pierces the host cell membrane, allowing the passage of infectious sporoplasm into the host cell cytoplasm. If a spore is phagocytosed by a host cell, germination will occur and the polar tube can pierce the phagocytic vacuole, delivering the sporoplasm into the host cell cytoplasm [1]. In XID APerC, we observed the extrusion of the polar filaments of intracellular mature spores from the phagolysosomes at 1 h, suggesting that spores may germinate after phagocytosis and escape. This extrusion of the polar tubule has been described in the literature, but ultra-micrographic analysis showing this process was not performed before. Therefore, this image is unprecedented in the literature. These findings indicate an intimate relationship between spores and the phagocytic vacuole, suggesting that some XID APerC macrophages have less microbicidal activity. We hypothesize that these macrophages can be polarized to M-2 profile in the absence of B-1 cells and promote the maintenance of *E. cuniculi*.

Through the secretion of IL-10, B-1 cells decrease NO production by macrophages, a phenomenon that may be associated with the lower microbicidal activity [12]. In this study, IL-10 production in infected and uninfected BALB/c APerC was not different, which perhaps explains why the macrophages of these cultures did not show increased NO production despite marked microbicidal activity. NO production by macrophages plays an important role in the elimination of intracellular parasites [42] and the literature has demonstrated that reactive nitrogen intermediates contribute to killing of *E. cuniculi* of peritoneal macrophages in mice by activated IFN-γ and LPS [43,44]. On the other hand, Franzen et al. [45] demonstrated that viable microsporidian spores did not induce the NO response in monocyte-derived human macrophages and that the number of intracellular spores and the amount of NO were negatively correlated. The authors suggested that modulation of the NO response by intracellular microsporidia may contribute to their survival within the macrophage, by an unknown mechanism. The role of NO in immunity against *E. cuniculi* was dismissed by Khan and Moretto [46] after evaluating its importance in protection against *E. cuniculi* infection using iNOS^-/-^ mice. The authors reported that none of the animals died or exhibited any signs or symptoms of disease throughout the course of the experiment and appeared clinically similar to the wild-type controls. The induction of NO in macrophages requires TNF, which is an important cytokine in the defense against intracellular pathogens as it promotes phagocytosis and intracellular killing [45]. In this study, there were no significant changes in levels of TNF and no difference in NO production between infected and non-infected cultures after 30 min, 1 h, and 48 h in all groups was observed. This phenomenon suggests that microbicidal activity was probably related to other mechanisms and should be investigated further.

B-1 cells from the peritoneal cavity of mice or cultures of adherent peritoneal cells can be clearly identified on the basis of their distinct morphology and cell surface phenotype. The main morphological characteristic of these cells is the formation of bridges of the nuclear membrane, suggesting a lobular organization of the nucleus. In addition, B-1 cells are characterized by small membrane projections and a large number of ribosomes, a predominance of euchromatin in B-1 cell nuclei, and more condensed chromatin in the nuclear periphery [21]. In this study, we identified B-1 cells by TEM in BALB/c APerC and B-1 CDP cultures. The B-1 CDPs decrease the expression of immunoglobulin M (IgM) but retain the expression of heavy-chain gene-variable VH11 or VH12, an immunoglobulin gene rearrangement that is predominantly expressed by B-1 cells [23]. The maintenance of lymphoid characteristics in B-1 CDPs is the characteristic of a unique type of phagocyte that is not related to monocyte-derived macrophages.

Cell-to-cell communication is required to guarantee proper coordination among different cell types within tissues. Studies have suggested that cells may also communicate by circular membrane fragments called micro vesicles that are released from the endosomal compartment as exosomes or shed off from the surface membranes of most cell types [47]. In this study, the identified phagocytic cells of B-1 CDP cultures had abundant micro vesicles in their membranes, indicating cell communication. In addition, we observed cells in the process of communicating by cell membrane projections (pseudopodia) or adhered cell-to-cell membrane between different types of cells in BALB/c APerC.

Th2 cytokines have been demonstrated in *E. cuniculi* infection [15,46] and an increase in the mRNA for IL-10 was observed in the splenocytes of infected animals [46]. This cytokine has been reported to be involved in the regulation of Th1 immune response against *Toxoplasma gondii* [48] and possibly has a similar role in *E. cuniculi* infection. In an *in vitro* study on cell cultures of human macrophages incubated with *E. cuniculi* spores, an increase in the levels of IL-10 was observed in supernatants of infected cultures, whereas this cytokine was below the limit of detection in the control group (non-infected cultures). Our findings demonstrated an increase in levels of IL-10 in the infected group of XID mice after 30 min and 1 h of infection, although this correlation was not statistically significant.

Chemokines are a group of small molecules that regulate cell trafficking of leukocytes. They mainly act on monocytes, lymphocytes, neutrophils, and eosinophils, and play an important role in host defense mechanisms [49]. MCP-1, also known as CCL2, was the first human chemokine to be characterized [50]. This molecule attracts cells of the monocyte lineage, including macrophages, monocytes, and microglia [51]. Mice deficient in MCP-1 were reported to be incapable of effectively recruiting monocytes in response to an inflammatory stimulus, despite the presence of normal numbers of circulating leukocytes [52]. Chemokine production has been documented to be induced by microsporidian infections in human macrophages. A primary human macrophage culture from peripheral blood mononuclear cells was infected with *E. cuniculi*, revealing the involvement of several chemokines, including MCP-1, in the inflammatory responses [53]. Our findings demonstrated an increase in the levels of MCP-1 in the supernatants of infected XID APerC after 30 min, 1 h, and 48 h of infection, although this correlation was not statistically significant. This cytokine was not detected after 96 h of infection.

IL-6 is a proinflammatory cytokine. In a previous study, we demonstrated that diabetes mellitus (DM) increased the susceptibility of C57BL/6 mice to encephalitozoonosis and DM mice infected with *E. cuniculi* showed higher levels of IL-6 than DM-*uninfected* mice, suggesting that DM may also modulate a pro-inflammatory state of the organism [54]. In the current study, we observed an increase in IL-6- in XID APerC after 30 min, 1 h, and 48 h of cultures infection. In contrast, the levels of this cytokine increased after 96 h and 144 h of infection in BALB/c APerC.

With the results obtained herein, we have demonstrated that B-1 cells modulate the activity of peritoneal macrophages infected with *E. cuniculi* to an M1 profile. Furthermore, part of these cells become B-1 CDPs having microbicidal activity against the pathogen, which explains the lower susceptibility of BALB/c mice to encephalitozoonosis associated with innate immune response.

## Methods

### Study Animals and Ethics Statement

Inbred specific pathogen free (SPF) BALB/c and BALB/c XID female mice at 6–8 weeks of age were obtained from the animal facility at Centro de Desenvolvimento de Modelos Experimentais (CEDEME), UNIFESP, Brazil. The animals were housed in polypropylene microisolator cages with a 12-hour light-dark cycle, maintained at 21 ±2 °C and >40% humidity, and fed on standard chow and water *ad libitum*.

### Ethics Statement

All the experimental procedures were performed in accordance with guidelines of Conselho Nacional de Controle de Experimentação Animal (CONCEA) and were approved by the Ethics Committee for Animal Research at Paulista University, under protocol number 385/15.

### *E. cuniculi* Spores

Spores of *E. cuniculi* (genotype I) (from Waterborne Inc., New Orleans, LA, USA) that were used in this experiment were previously cultivated in a rabbit kidney cell lineage (RK-13, ATCC CCL-37) in Eagle medium supplemented with 10% of fetal calf serum (FCS) (Cultilab, Campinas, SP, Brazil), pyruvate, nonessential amino acids, and gentamicin at 37 °C in 5% CO_2_. The spores were purified by centrifugation and cellular debris was excluded by 50% Percoll (Pharmacia) as described previously [55].

### Adherent Peritoneal Cells (APerC) and B-1 Cell-Derived Phagocyte (B-1 CDP) Cultures

The APerC were obtained from the peritoneal cavities (PerC) of BALB/c and B-1 cell-deficient XID mice. PerC were washed using RPMI-1640 medium and 0.5×10^6^ cells were dispensed in each well of 24-well plates and incubated at 37 °C in 5% CO_2_ for 40 min. Non-adherent cells were discarded and RPMI-1640 supplemented with 10% FCS (R10) was added to the adherent fraction. To obtain B-1 CDP, APerC from BALB/c mice were cultured and the enriched B-1 cells in the floating medium were collected from the third day [56]. Cultures of 0.5 × 10^6^ cells/well were re-suspended in R10 and re-cultured under the same conditions as described above.

### *E. cuniculi* Infection

The culture of BALB/c APerC, XID APerC, and B-1 CDP were infected simultaneously with *E. cuniculi* (1×10^6^ spores/mL) in the proportion of two spores per cell (2:1). The cultures were incubated at 37 °C in 5% CO_2_ for 30 min, 1h, 48 h, 96 h, and 144 h after infection, following which the supernatants were collected and stored at –80 °C to measure the level of NO and cytokines. The cultures uninfected with *E. cuniculi* were incubated for the same time intervals and used as a group control.

### Measurement of NO Production

The production of NO was measured using a colorimetric indirect method based on the detection of nitrite (nitrate was initially converted to nitrite by nitrate reductase) as a product of the Griess reaction (R&D Systems) in the supernatants of cell cultures. Briefly, supernatants were mixed with the Griess reagent [equal volumes of 0.2% (w/v) naphthylethylenediamine in 60% acetic acid and 2% (w/v) sulfanilamide in 30% (v/v) acetic acid] and incubated at room temperature for 10 min. A spectrophotometer was used to measure the absorbance at 540 nm and fresh culture medium was used as a blank in all the experiments. The amount of nitrite in the test samples was calculated from a sodium nitrite standard curve (0.78–100 µM).

### Quantification of Cytokines

Cytokines were measured in culture supernatants using Cytometric Bead Array (CBA) Mouse Inflammation Kit (BD Bioscience, San Jose, CA, USA). The kit was used for the simultaneous detection of mouse monocyte chemoattractant protein-1 (MCP-1), interleukin-4 (IL-4), IL-6, IFN-γ, tumor necrosis factor (TNF-α), IL-10, and IL-12p70, according to the manufacturer’s instructions. Briefly, the supernatant samples were added to bind to allophycocyanin (APC)-conjugated beads specific for the cytokines listed above and phycoerythrin (PE)-conjugated antibodies. The samples were incubated for 2 h at room temperature in the dark, then measured using FACS Canto II Flow Cytometer, and analyzed by FCAP Array**^TM^** Software (BD Bioscience), version 3.0. Individual cytokine concentrations (pg/mL) were indicated by the intensity of PE fluorescence and cytokine standard curves.

### Flow Cytometry of Peritoneal Cells

Infected and uninfected APerC were detached using a cell scraper and washed with PBS. The cell suspensions were centrifuged and subsequently washed with PBS and re-suspended in 100 μL PBS supplemented with 1% bovine serum albumin (BSA) (PBS-BSA 1%). Each sample was incubated at 4°C for 20 min with anti-CD16/CD32 to block the Fc II and III receptors. After incubation, the cells were washed, divided into two aliquots, and re-suspended in PBS-BSA 1%. Each sample was then incubated with the following monoclonal antibodies: **1)** fluorescein-isothiocyanate (FITC)-conjugated rat anti-mouse CD80/CD86, 2) PE-Cyanine 5 (PE-Cy5)-conjugated anti-mouse CD40, 3) APC-Cy7-conjugated rat anti-mouse CD11b, and 4) Alexa Fluor*®* 647 rat anti-mouse CD206 (BD-Pharmingen, San Diego, CA, USA) for analysis of the surface markers. The cell pellet was incubated with fluorochrome-conjugated antibodies for 20 min at 4°C, washed with PBS-BSA 1%, re-suspended in 500 μL of PBS, and analyzed with BD Accuri C6 flow cytometer.

### Phagocytic Capacity and Index

To investigate the phagocytic capacity and index, 10 μL Calcofluor (Sigma-Aldrich, St. Louis, USA) was added per milliliter of the cell cultures to visualize the spores inside the phagocytic cells. Phagocytic capacity and index were calculated according to the formula: FC = number of phagocytes containing at least one ingested spore/100 phagocytes and FI = total number of phagocytic spores/100 phagocytes containing spores.

### Measurement of dead cells

Cell cultures were washed twice with cold PBS and then resuspended in 1x annexin-binding buffer (BD Biosciences) at a concentration of 1 × 10^6^ cells/mL. Thereafter, 100 μL of the solution (1 × 10^5^ cells) was transferred to a 5-mL culture tube, to which 1 μL of PE Annexin V and 1 μL of 7-AAD were added and followed by incubation for 15 min on ice in dark conditions. Subsequently, 400 μL of 1x annexin binding buffer was added to the samples, incubated for 30 min, and all the resultant cell suspensions were analyzed using the BD Accuri C6 flow cytometer.

### Analyses using Light and Transmission Electron Microscopy

The cell volume of BALB/c APerC, XID APerC, and B-1 CDP cells, cultured as described above, was adjusted to 1×10^7^ cells. Each culture was transferred to 25 cm^2^ bottles and incubated in the same medium containing 10% FCS (R10) at 37 °C with 5% CO_2_ for 40 min. The culture medium was then removed and fresh R10 medium containing *E. cuniculi* spores (2:1) was introduced into the bottles. The cultures were collected after 1 h, 48 h, and 144 h of incubation and fixed using 2% glutaraldehyde in 0.2 M cacodylate buffer (pH 7.2) at 4 °C for 10 h. They were then post-fixed in 1% OsO_4_ buffer for 2 h. Semi-thin sections stained with toluidine blue were made for visualization by light microscope, and ultrathin sections were made for TEM analysis.

### Statistical Analysis

The groups were compared using the two-way analysis of variance (ANOVA) and the significance of the mean difference within and between the groups was evaluated by multiple comparisons using the Bonferroni or Tukey’s post-tests. All the experimental data were expressed as mean ± standard error mean, indicated by bars in the figures. *P* values <0.05 were considered statistically significant. All the graphs and statistical analyses were made using GraphPad Prism software version 6.0 for Windows (GraphPad Software, San Diego, California, USA).

## ACKNOWLEDGMENTS

We acknowledge Fundação de Amparo à Ciência do Estado de São Paulo-Fapesp and Capes for financial support.

## Supporting Information Legends

Balb and Xid adherent peritoneal cells were cultured in vitro and challenged with *E. cuniculi*. Balb macrophages had a predominant M1 profile, while those of Xid had a more M2 profile

